# Energy Conservation as a Measure of Simulation Accuracy

**DOI:** 10.1101/083055

**Authors:** Peter Eastman, Vijay S. Pande

## Abstract

Energy conservation is widely used as a measure of accuracy for molecular simulations. When reporting rates of energy drift, researchers usually assume it is linear in the simulation length, temperature, and system size. We study these assumptions and find that all three are incorrect. Energy drift is too complicated to characterize with a single number, and a more sophisticated analysis is needed to identify the effects of systematic versus random drift, and of integration error versus numerical error. We further argue that energy conservation is not a reliable measure of accuracy. Having small overall drift on long time scales is not a sufficient condition, and in some cases not a necessary condition, for a simulation to produce meaningful results.

## 1. Introduction

Energy conservation is widely used as a way to assess the accuracy of molecular simulations. The idea is that, if all sources of error have been sufficiently minimized, there should be negligible change in the total energy of a system over the course of a simulation. Measuring how much the energy actually changes therefore serves as a test of how much error is present in the simulation.

We emphasize that our concern is primarily with energy conservation used *as a measure of simulation accuracy*. There are many situations where energy conservation is important for its own sake, but those are not the focus of this paper. The majority of molecular simulations are performed at constant temperature, so they do not even attempt to conserve energy. However, researchers often use energy conservation in the absence of a thermostat to assess the accuracy of their production simulations, even if the production simulations do include a thermostat. That is the subject we wish to study.

Given its importance and prevalence in the literature, it is surprising how little agreement there is on how to measure energy conservation. We have reviewed a number of papers that report energy drift values for different simulation codes, and find an enormous variation in their methods.[1-5] They differ widely in the systems they simulate, the trajectory lengths, and the methods used to calculate energy drift.

For example, the simulation lengths reported in these papers cover two orders of magnitude, from 1 ns up to 100 ns. They universally normalize the energy drift by simulation length, reporting it in units of energy per time. This normalization assumes that energy drift is linear in time. Indeed, most authors appear to explicitly assume this. Results from simulations of different lengths are often presented side by side in a single table, indicating to the reader they can be directly compared. In some cases,[1, 2] the simulation length is never even stated; it is simply assumed that the normalization makes this information irrelevant.

Surprisingly, there seems to have been very little discussion of this assumption in the literature. We are aware of no justification, either theoretical or empirical, that has been offered for why it should be expected to hold. On the contrary, some theoretical work has suggested energy drift should be primarily diffusive rather than linear in time.[6, 7] But if this assumption is wrong, then results from simulations of different length are not equivalent and cannot be directly compared to each other.

Similar considerations apply to system size and temperature. The systems reported in these papers cover a huge range, from as few as 304 atoms to as many as 92,224. Temperatures of 300K and 394K are used in different cases, while some papers[2, 4] do not even report the temperatures of their simulations. In all cases, the results are normalized by temperature and number of degrees of freedom, or equivalently, by the average kinetic energy. This normalization assumes that energy drift should be linear in temperature and number of degrees of freedom, and once again, this assumption seems to be explicitly made by most authors. For example, results for systems of different size are often presented side by side and directly compared to each other. But here again, we are aware of no justification for why these assumptions should be expected to hold.

Even the simple question of how to compute a single “energy drift” value from simulation data is usually left unspecified.[1, 2, 4, 5] Most authors appear to have felt this was obvious and did not need to be stated, yet one can easily think of several different ways to do it:

1. Compute the energy difference between the first and last frames in the trajectory, then divide by the length of the trajectory.

2. Perform a linear regression to fit a straight line to the curve of energy vs. time and report its slope.

3. Perform either 1 or 2 for multiple simulation trajectories, then report the mean energy drift across all trajectories; or the mean absolute value of the energy drift; or the root-mean-squared energy drift.

Clearly these methods will produce different results, and values calculated with different methods cannot be directly compared to each other.

We encountered these issues while benchmarking OpenMM.[8] Our goal was to study its accuracy when using different program options and simulation parameters, then compare to the published numbers for other simulation codes. We soon realized the impossibility of that comparison, which led us to conduct a deeper study of the nature and significance of energy drift. All simulations described in this paper were conducted with OpenMM, but most of the conclusions are generally applicable to a wide range of molecular simulation codes.

The rest of this paper is organized as follows. In sections 2 and 3 we consider various issues involved in measuring and analyzing energy drift. Sections 4, 5, and 6 consider each of the assumptions (linearity in time, temperature, and system size) in turn to assess their validity. In section 7, we turn to an even more fundamental question: to what extent is energy conservation actually a useful measure for evaluating simulation accuracy? The following are our major conclusions:

1. All of the assumptions discussed above are incorrect. Therefore, energy drift values calculated for different sized systems, from different length trajectories, or at different temperatures cannot be directly compared to each other.

2. Energy conservation is too complicated to fully characterize with a single number. A more detailed analysis is required to measure it in a meaningful way.

3. Energy conservation is a singularly bad proxy for accuracy. Just because a simulation has low energy drift on long time scales, one cannot conclude the equations of motion are being integrated accurately.

## 2. Computation of Energy

How should one compute the energy at a given instant in a simulation? This seemingly trivial question turns out to be somewhat subtle, and is often done in unnecessarily inaccurate ways.

Most constant-energy molecular simulations use some variation on the Verlet integrator:[9]

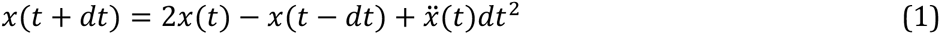

where *x* is the vector of particle coordinates, *t* is the current time, and *dt* is the step size. There are two important things to note about this integration method. First, it has a local error that scales as O(*dt*^4^) and a global error that scales as O(*dt*^2^). Second, it is formulated entirely in terms of the positions. No velocities appear explicitly in it. This is important, because we must somehow determine velocities in order to calculate the kinetic (and hence total) energy.

The Verlet integrator is commonly reformulated in one of two ways to introduce a velocity-like variable into it. The “velocity Verlet” algorithm defines

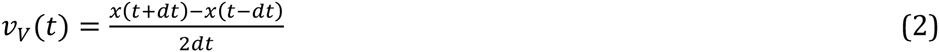

Equation 1 can then be rewritten as a series of three steps:

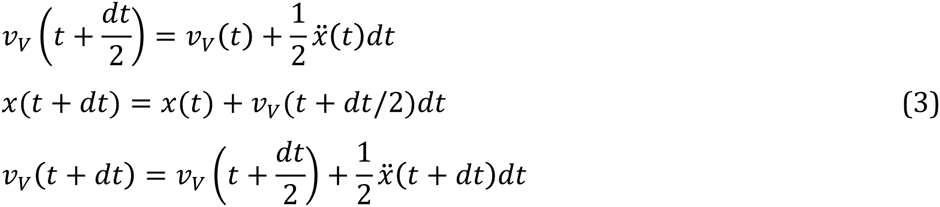

*vV*(*t*) is a finite difference approximation for velocity that is accurate to second order. It is best not to think of it actually being the velocity, however. It is an internal parameter of the integrator that is exactly defined by equation 2. It happens to equal a low order approximation to the velocity, but the trajectory (and hence the “true” velocity) has a local error of O(*dt*^4^), not O(*dt*^2^).

The “leapfrog Verlet” algorithm defines

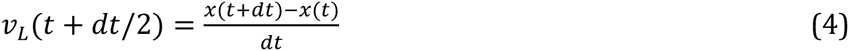

Equation 1 can then be rewritten as

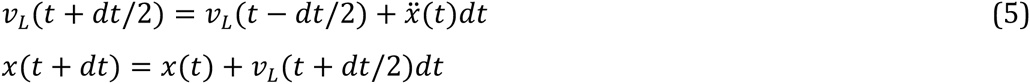

*v_L_*(*t*+*dt*/2) is a second order approximation to the velocity at time *t*+*dt*/2. It also is a first order approximation to the velocity at time *t*. As with velocity Verlet, however, these identifications can easily be misleading. *v_L_* is simply an internal parameter of the algorithm that is exactly defined by equation 4.

Higher order approximations to the velocity can be formed by combining the positions at more points in time.[10] The “five point stencil” is given by:

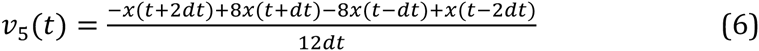

This is accurate to fourth order, the same as the local error in the Verlet algorithm. The integration is still performed with one of the Verlet integrators given above, but equation 6 is used in place of *v_V_* or *v_L_* when computing the kinetic energy. Still higher order approximations exist, but there is no value in using them: because the positions are only accurate to fourth order, no combination of them can produce a velocity more accurate than that. We therefore recommend equation 6 for calculating velocities, and hence energies. It is used throughout this paper.

## 3. Types of Error

The errors discussed in the previous section are inherent in the integration method. They come from using a finite sized time step, and are present even when all calculations are done to infinite precision. We refer to this as “integration error”. (It is also sometimes referred to as “truncation error”, since it is caused by omitting higher order terms in the integration method.)

In practice, the calculations are never done to infinite precision. This introduces other types of errors into the simulation, which we refer to as “numerical error”. Many types of approximations can lead to numerical error. Some common examples include:

1. Numeric representations that only use a finite number of bits of precision.

2. Cutoffs on nonbonded interactions (in real space, and possibly also in reciprocal space) that create discontinuities in the energy.

3. Neighbor lists that are not updated every time step, allowing interactions to occasionally be missed.

4. Lookup tables used to approximate expensive functions.

These sources of error are very different in their origins, but for simplicity we group them all into the single category of “numerical error”. They are all ways in which the integration algorithm is not computed exactly, to infinite precision. They therefore introduce error into the trajectory, above and beyond the error that is inherent in the integration algorithm.

There are some important differences between integration error and numerical error. The magnitude of integration error depends strongly on the time step: the global error scales as O(*dt*^2^) for all Verlet integrators. Numerical error usually scales as a lower power: it may be independent of *dt*, or in some cases actually increase with decreasing step size. For example, if an approximation introduces a constant level of error in every time step, the total error during a time interval will scale with the total number of time steps, or 1/*dt*. This means that for sufficiently large step size, numerical error is negligible compared to integration error, while for sufficiently small step size, integration error is negligible compared to numerical error.

Another important difference is that, because integration error is inherent in the Verlet algorithm, it is identical for all simulation codes so long as they use the same step size and simulate the same system. Numerical error, on the other hand, depends very sensitively on the implementation details. It comes from the fact that forces are not computed exactly and the integration algorithm is not performed to infinite precision. This means that our measurements of integration error in the following sections should be universal and widely applicable, whereas our measurements of numerical error are specific to OpenMM.

We also distinguish between systematic errors and random errors. Systematic errors are the same for every time step, whereas random errors are uncorrelated between time steps (at least if those steps are sufficiently separated in time). They lead to very different behaviors. Systematic errors add together, leading to a linear change in energy with time, while random errors produce a diffusive drift whose magnitude grows as the square root of time.

It is tempting to conclude that systematic errors are “more important” than random ones, since they produce a much larger drift over sufficiently long time periods. That conclusion is not justified however. Random errors are still errors, no less for being uncorrelated, and they can still affect simulation results.

It is important to remember that many molecular simulations are performed at constant temperature, not constant energy. When using a stochastic thermostat, errors cannot produce an energy drift over long time periods, because it is counteracted by the thermostat. In this case, when one measures the energy drift in a constant energy simulation, the primary goal is to show that the equations of motion are being integrated accurately, and numerical and integration errors are not cause for concern. A large random error clearly shows that they are *not* being integrated correctly, and the accuracy of the simulation *cannot* be assumed.

The question of exactly how errors impact one’s results is beyond the scope of this article. It might depend in complicated ways on the type of errors, the type of results one is interested in, and many other details of the simulation. A particular type of error might have a large effect on some results, but no effect at all on other results. Regardless of these details, all results must be treated as suspect if the equations of motion are not being integrated accurately. They might still be correct, but that cannot be assumed until the researcher presents convincing evidence. Systematic and random energy drift are both signs of inaccurate integration. Any analysis of energy conservation therefore needs to consider both systematic and random errors and evaluate their importance independently.

The above discussion applies to system size as well as to simulation length. Errors may be the same for every degree of freedom, or uncorrelated between degrees of freedom. The former case produces a total error in the energy that scales with the number of degrees of freedom as O(*n*), while the latter (being a sum of uncorrelated random variables) produces a total error that scales as O(√*n*).

## 4. Scaling With Simulation Length

We now turn to the first widely made assumption: that energy drift is linear in the simulation length. Figure 1 shows the energy as a function of time for a 5 ns simulation of ubiquitin[11], a 1231 atom protein in GBSA-OBC implicit solvent[12]. This simulation used the Amber 99SB force field[13], a 1 fs time step, a leapfrog Verlet integrator, and no cutoff on the nonbonded interactions. No degrees of freedom were constrained. (The choice to use implicit solvent with no cutoff was made to eliminate any source of error coming from nonbonded cutoffs, and thus let us more clearly see the effect of other types of error. When using PME, the error depends in a complicated way on many different parameters, and that would unnecessarily complicate the analysis.) The energy was recorded every 1 ps.

**Figure 1.**
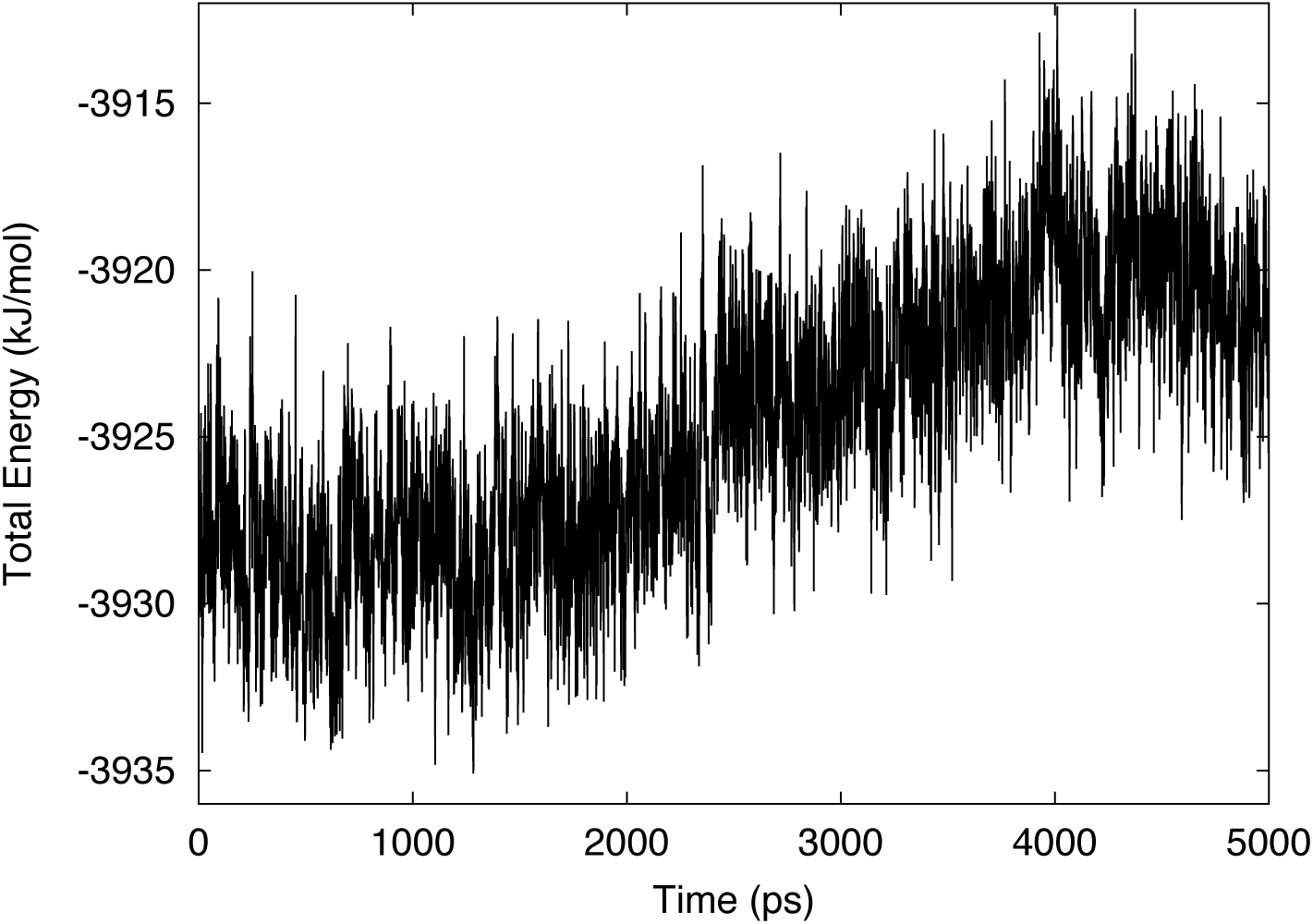
Total energy versus time for a 5 ns simulation of ubiquitin in implicit solvent. This simulation used double precision and a 1 fs time step.

It is immediately clear that the energy drift is primarily diffusive, not linear. It fluctuates irregularly on both short and long time scales, but the net effect is to change very little over the course of the simulation. This is our first indication that the assumption of linear drift is wrong.

To make the analysis more quantitative, we compute the difference between each adjacent pair of energies. This yields 5000 samples of how much the energy changes in 1 ps. A histogram is shown in Figure 2. It clearly resembles a symmetric distribution whose center is very close to 0.

**Figure 2.**
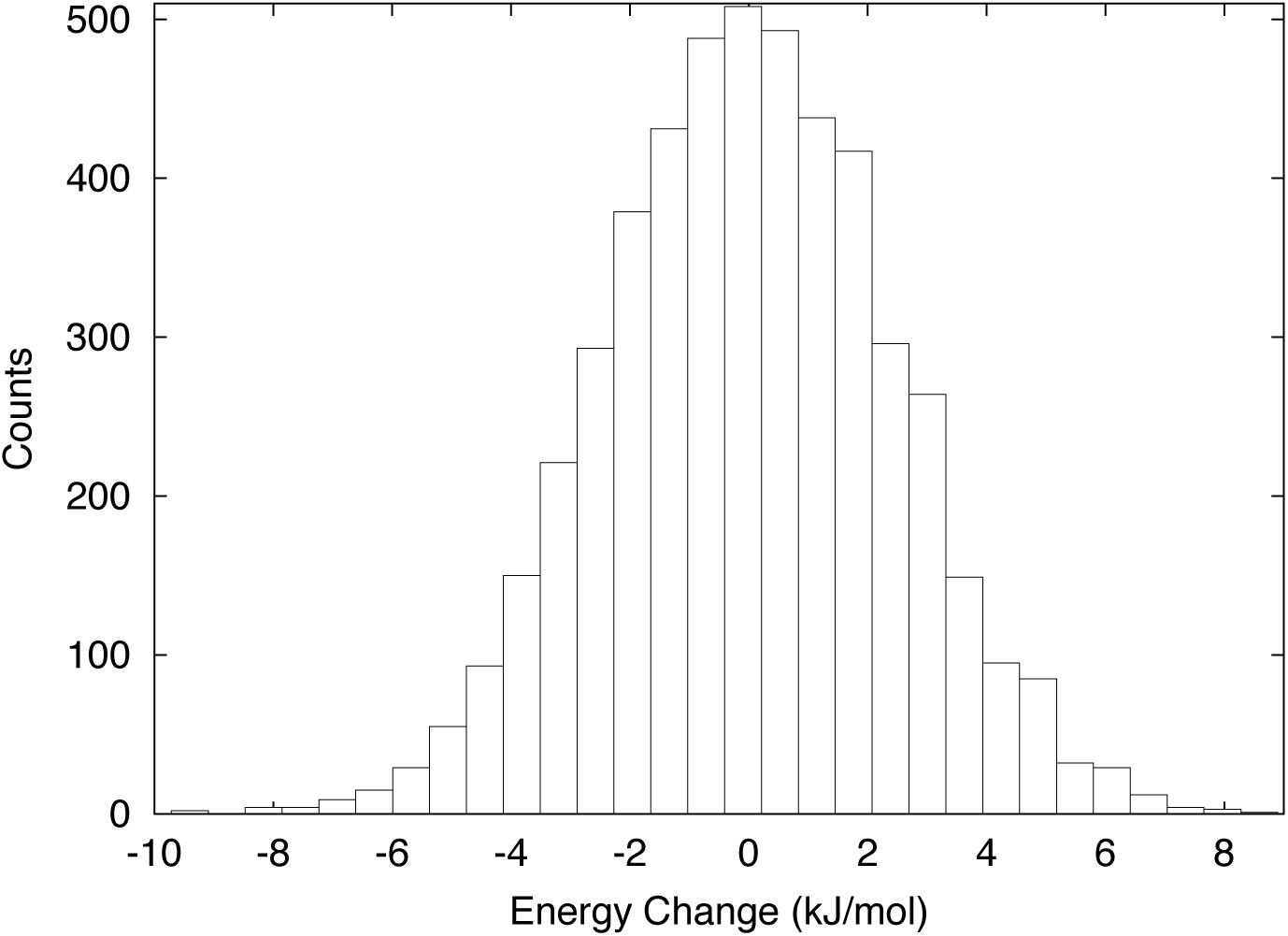
Histogram of the change in total energy over 1 ps intervals for the simulation shown in Figure 1.

Statistics for the distribution are given in Table 1. We repeated the simulation with time steps of 1.0, 0.5, and 0.25 fs. OpenMM allows calculations to be done in three different precision modes: nearly all single precision; all double precision; and a mixed precision mode that computes forces in single precision but does integration in double precision. We therefore ran simulations in all three modes. For each one, we report the mean (which gives a measure of the systematic error), standard deviation (which gives a measure of the random error), and standard error (which reflects the uncertainty in our measurement of the mean). Note that the mean is simply equal to the total change in energy over the course of the simulation, divided by the number of 1 ps intervals making up the simulation. A more robust measure of systematic error is to perform a linear regression to the curve of energy versus time. The slope of that line, when measured in kJ/(mol·ps), should be directly comparable to the mean of the 1 ps energy change distribution. This value is also shown in Table 1.

The first thing to notice is that in all cases, the standard deviation is orders of magnitude larger than the mean. This means that for short simulations, the energy drift will be completely dominated by random error and its magnitude will scale as the square root of the simulation length. For sufficiently long simulations, on the other hand, even a tiny systematic error will eventually become dominant. In either case, simply dividing by the simulation length and reporting a single drift rate is misleading. If the simulations are short, all information about systematic error will be lost. If the simulations are long, all information about random error will be lost. In contrast, the analysis shown in Table 1 allows us to break them apart and measure each one independently.

**Table 1.** Statistics for a set of 5 ns simulations of ubiquitin in implicit solvent.. The mean, standard deviation, and standard error describe the distribution of energy changes over 1 ps intervals. The slope is from a linear regression to the curve of energy versus time for the whole simulation.

| Precision | Time Step<br>(fs) | 1 ps Energy Change Distribution |  |  | Slope<br>(kJ/(mol·ps)) |
| --- | --- | --- | --- | --- | --- |
|  |  | Mean<br>(kJ/(mol·ps)) | Standard Deviation<br>(kJ/(mol·ps)) | Standard Error<br>(kJ/(mol·ps)) |  |
| single | 1.0 | 0.00300 | 2.35 | 0.0332 | 0.00163 |
| single | 0.5 | 0.00373 | 0.647 | 0.00914 | 0.00398 |
| single | 0.25 | 0.00767 | 0.283 | 0.00400 | 0.00719 |
| mixed | 1.0 | 0.00144 | 2.23 | 0.0315 | 0.00119 |
| mixed | 0.5 | 0.000354 | 0.613 | 0.00867 | 0.000417 |
| mixed | 0.25 | -0.000249 | 0.158 | 0.00224 | -0.000124 |
| double | 1.0 | 0.00191 | 2.43 | 0.0343 | 0.00226 |
| double | 0.5 | -0.000231 | 0.636 | 0.00900 | 0.00000100 |
| double | 0.25 | -0.0000605 | 0.169 | 0.00239 | -0.0000000710 |

We can estimate how well converged these values are by splitting each trajectory in half, then computing the statistics independently for the first half and the second half. In all cases, this produces two values for the standard deviation that agree with each other to within a few percent. We conclude that the measures of random error in Table 1 are well converged, and the simulations are long enough to measure it accurately.

The measures of systematic error, on the other hand, are not well converged. There often are significant differences between the estimates from the two halves of a trajectory. This should not be surprising, given the much larger magnitude of the random error. On any time scale short enough for the random error to be significant, the rate of energy drift is not deterministic. The total drift in any one simulation is just a sample from a distribution. Performing several independent simulations will produce a different drift for every one. It is therefore essential that when one reports a “drift rate” for simulations of a given length, the simulation be repeated several times and a distribution be reported instead of a single value.

To illustrate this fact, we performed additional simulations of three different lengths: 10, 100, and 1000 ps. For each length, 10 independent simulations were performed. Before starting a simulation, the system was first equilibrated at 300K, and the simulations differed only in the random number seed used for the equilibration. All simulations used single precision and a 1 fs time step.

For each simulation, we performed a linear regression to the curve of energy versus time. This yielded 10 independent “drift rates” for each simulation length, allowing us to study the distribution of values. Table 2 shows the mean, minimum, and maximum values for each simulation length, along with their standard deviation. In all cases, the width of the distribution is larger than its mean, emphasizing the fact that any one value taken on its own is meaningless. The width of the distribution decreases as the simulation length increases, but even at 1 ns the values are clearly still dominated by random error.

**Table 2.**
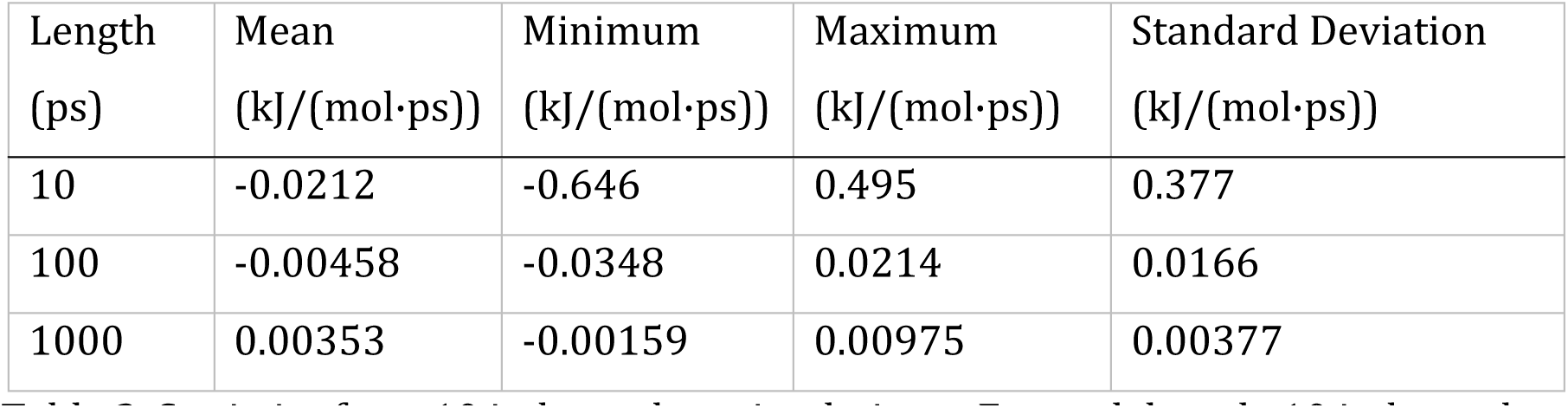
Statistics from 10 independent simulations. For each length, 10 independent simulations were run and a linear regression to energy versus time was performed. This yields 10 independent “drift rates” whose distribution is characterized by the statistics shown.

| Length<br>(ps) | Mean<br>(kJ/(mol·ps)) | Minimum<br>(kJ/(mol·ps)) | Maximum<br>(kJ/(mol·ps)) | Standard Deviation<br>(kJ/(mol·ps)) |
| --- | --- | --- | --- | --- |
| 10 | -0.0212 | -0.646 | 0.495 | 0.377 |
| 100 | -0.00458 | -0.0348 | 0.0214 | 0.0166 |
| 1000 | 0.00353 | -0.00159 | 0.00975 | 0.00377 |

Returning to Table 1, we next compare the mean to the standard error, which reflects the uncertainty in our measurement of the mean. We immediately notice something odd: in all of the mixed and double precision cases, as well as single precision with a 1 fs time step, the mean is at least an order of magnitude smaller than the standard error. This is surprising. If the data really consisted of independent samples from a normal distribution, we would expect the mean to be similar to the standard error, since it is only being measured to that level of accuracy. Instead we find that in the majority of cases, it is much smaller than we have any right to expect.

This is not a coincidence. It reflects the fact that Verlet integrators are symplectic, and hence conserve energy over long time scales much better than one would expect given the magnitude of integration error. We will discuss this issue further in section 7.

The above assumes the samples really are independent. That assumption is not guaranteed to be true. On sufficiently short time scales, we expect the changes in energy to contain correlations. This is clearly seen in Figure 3, which shows the evolution in energy over a very short piece of a simulation. It contains clear oscillations at high frequencies, reflecting the presence of oscillatory motions within the system being simulated. On any time scale short enough for the motion to contain correlations, it is possible for the changes in energy to also contain correlations.

**Figure 3.**
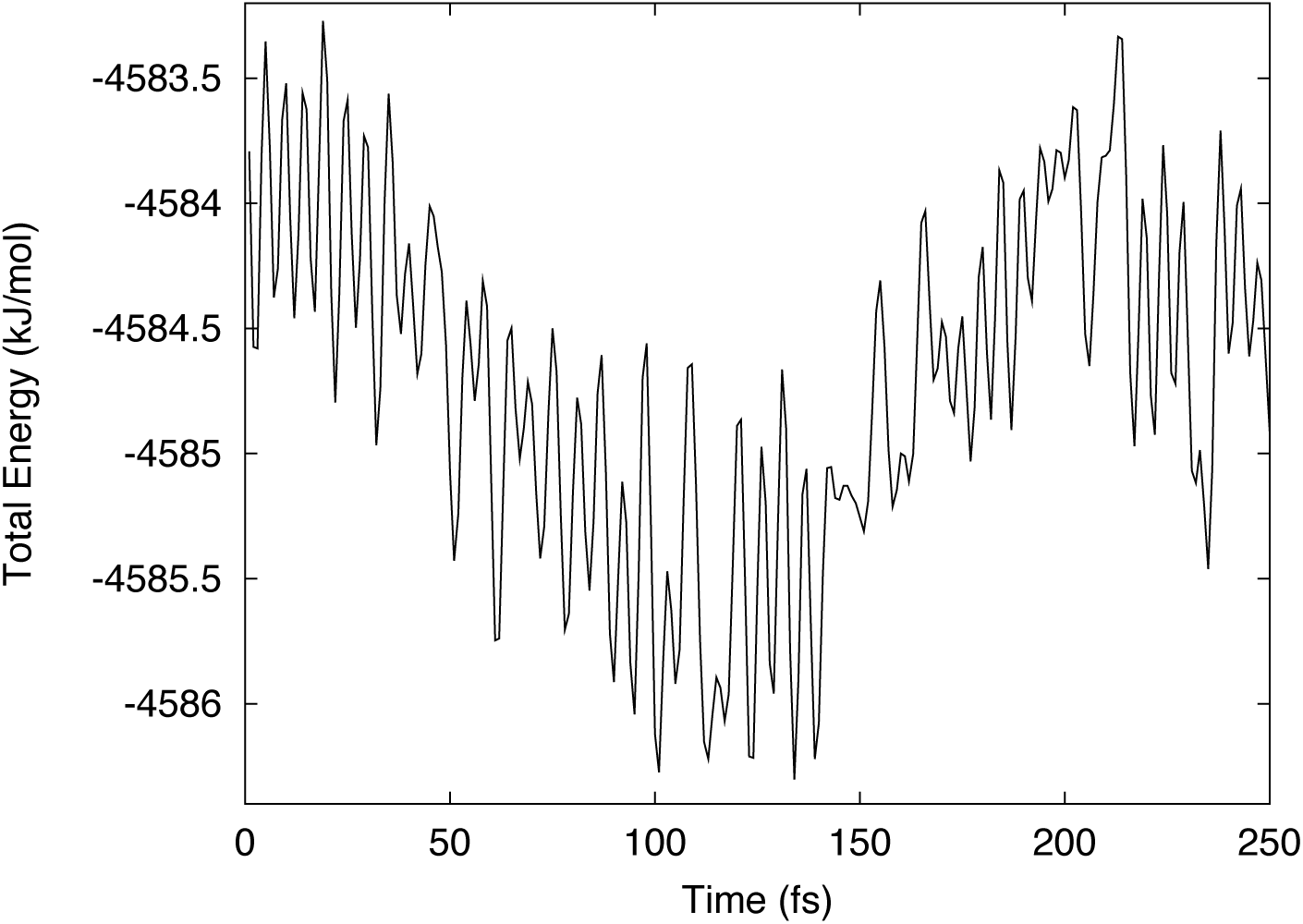
Total energy versus time for a short segment of a simulation of ubiquitin in implicit solvent. This simulation used double precision and a 1 fs time step.

An analysis of characteristic time scales in protein simulations shows that the lowest frequency oscillatory motions in proteins have periods less than 100 fs[14]. This fact motivated our choice of 1 ps as the sampling interval: it was chosen to be as short as possible, while still being much longer than the slowest oscillatory motions, and hence unlikely to contain significant correlations. Even if residual correlations are still present at 1 ps, this will have no effect on the measures of random and systematic drift calculated in Table 1. The only numbers that would be affected are the standard errors, which would be based on an overestimate of the number of independent measurements. They would therefore underestimate the true uncertainty in the mean, and the argument given above would then become even stronger.

Next consider how the error varies with step size, as shown in Figure 4. This lets us distinguish between numerical error and integration error, since the latter should scale as O(*dt*^2^), and hence should decrease by approximately a factor of 4 when the step size is cut in half. First we look at the double precision simulations. When the step size is decreased from 1 fs to 0.5 fs, the random error decreases by a factor of 3.8. When it is further decreased to 0.25 fs, the error decreases by another factor of 3.8. This is good agreement with the expected scaling, and suggests that even at a time step of 0.25 fs, the error primarily consists of integration error while numerical error makes little contribution to the energy drift. The results for mixed precision are similar: the random error decreases by 3.6 when going from 1 to 0.5 fs, and by 3.9 when going from 0.5 to 0.25 fs.

**Figure 4.**
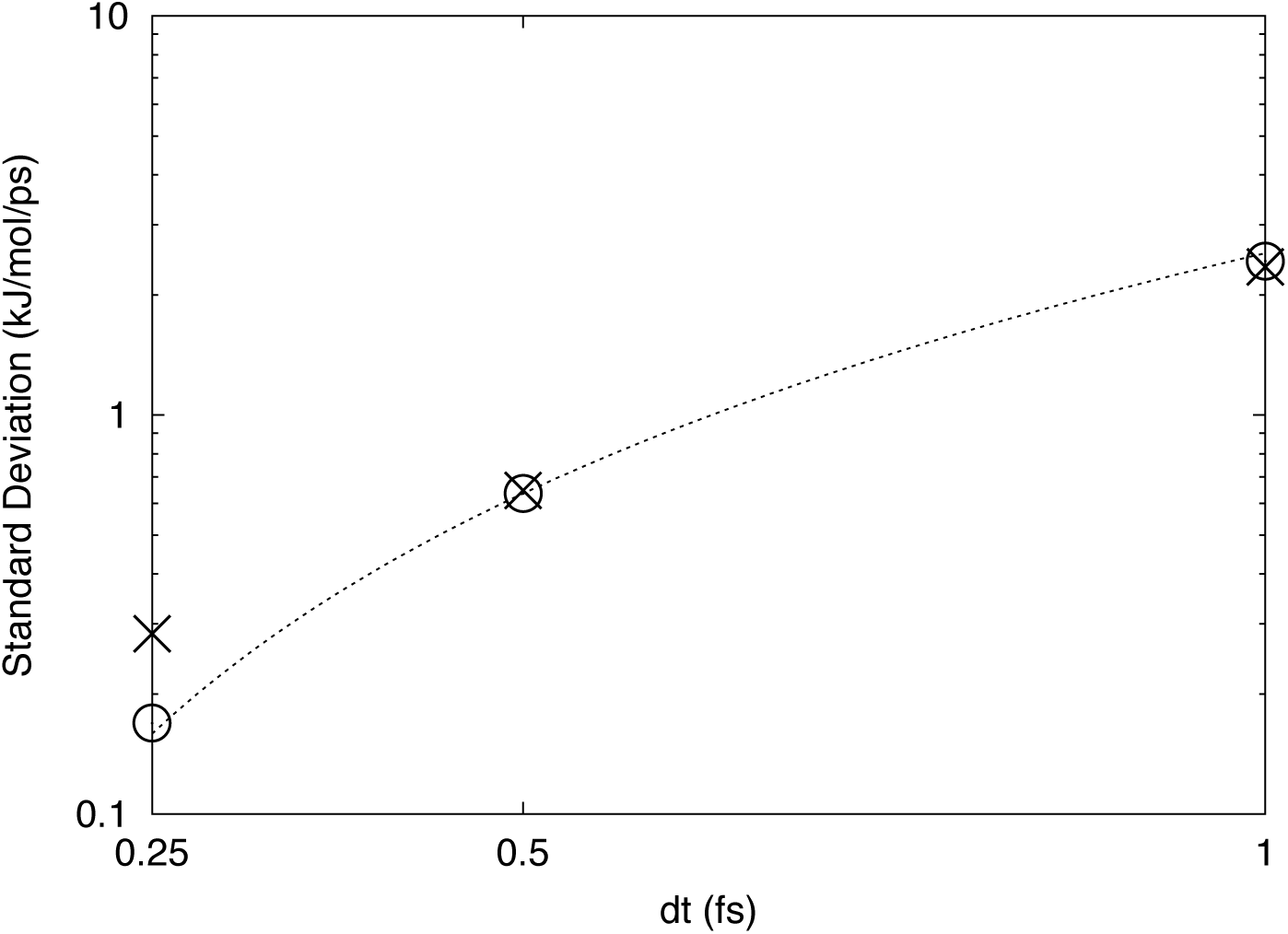
Standard deviation of the 1 ps energy change distribution versus step size. The values are for single precision (crosses) and double precision (circles). The dotted line is proportional to *dt*^2^ and is shown only for reference.

Now look at the single precision simulations. For 1 fs and 0.5 fs time steps, the random error is very close to the corresponding values for double precision. With a 0.25 fs time step, however, it decreases less than expected and is significantly larger than for double precision. This suggests that numerical error is starting to become significant. We also note that for both 0.5 fs and 0.25 fs time steps, the distribution mean is much larger than for double precision, and becomes comparable to the standard error. This is another indication that numerical error is significant, since the symplectic property of the integrator only affects the energy drift from integration error, not from numerical error.

## 5. Scaling With Temperature

Now we consider how energy drift varies with temperature. Some care is required in defining what we mean by this. When referring to the “temperature” of a simulation, one usually means that it is coupled to a heat bath by a thermostat, and one is specifying the temperature of that heat bath. A constant energy simulation is, by definition, not a constant temperature simulation. It is not coupled to a heat bath, and does not have a uniquely defined temperature.

Before starting a simulation, one usually first equilibrates it by simulating it at constant temperature for some amount of time. The temperature of that equilibration could be used as a measure of the “temperature” of the following constant energy simulation, but that is not reliable. The equilibration temperature does not guarantee anything about the energy of any one frame of the simulation: it merely sets a probability distribution for it. The last frame of the equilibration, which is then used as the first frame of the constant energy simulation, could easily happen to be an unusually high or low energy state for that temperature.

Another way to measure temperature is to compute the kinetic energy of the system. Since this is expected to have an average of kT/2 per degree of freedom, it can be used to compute an instantaneous effective temperature. Of course, it will not remain constant over the simulation, but as long as it does not fluctuate too much, the average instantaneous temperature over the simulation can serve as a reasonable measurement of the “temperature” of the simulation. This is what we do here.

The simulations described in section 4 were all equilibrated at 300K. To test the temperature dependence, we performed three additional simulations using mixed precision and a 1 fs time step. They were equilibrated at 100K, 200K, and 400K respectively. Table 3 lists the mean and standard deviation of the 1 ps energy change distribution for each one, along with the average temperature of the simulation and the slope of the linear fit to energy versus time.

**Table 3.** Statistics for a set of simulations of ubiquitin in implicit solvent at different temperatures. The mean and standard deviation describe the distribution of energy changes over 1 ps intervals. The slope is from a linear regression to the curve of energy versus time for the whole simulation. All simulations used mixed precision, a 1 fs time step, and 5 ns simulation length.

| Temperature<br>(K) | 1 ps Energy Change Distribution |  | Slope<br>(kJ/(mol·ps)) |
| --- | --- | --- | --- |
|  | Mean<br>(kJ/(mol·ps)) | Standard Deviation<br>(kJ/(mol·ps)) |  |
| 101.6 | 0.000768 | 0.676 | 0.000835 |
| 205.2 | 0.000279 | 1.45 | 0.000303 |
| 293.3 | 0.00144 | 2.23 | 0.00119 |
| 405.8 | 0.00109 | 3.19 | 0.00256 |

The standard deviation shows clear linear scaling with temperature, as shown in Figure 5. For short simulations that are dominated by random drift from integration error, the assumption that drift is proportional to temperature seems to be justified. On the other hand, the systematic error (as measured either by the mean or the slope) shows a very irregular variation with temperature, and is not even monotonic. For all simulations, the standard error (not shown) is much larger than the mean, so there is significant uncertainly is our estimates of the systematic error. It is possible that if we ran much longer simulations, a linear dependence on temperature would become apparent. But at the very least, our data does not support the assumption that energy drift is proportional to temperature for long simulations dominated by systematic drift.

**Figure 5.**
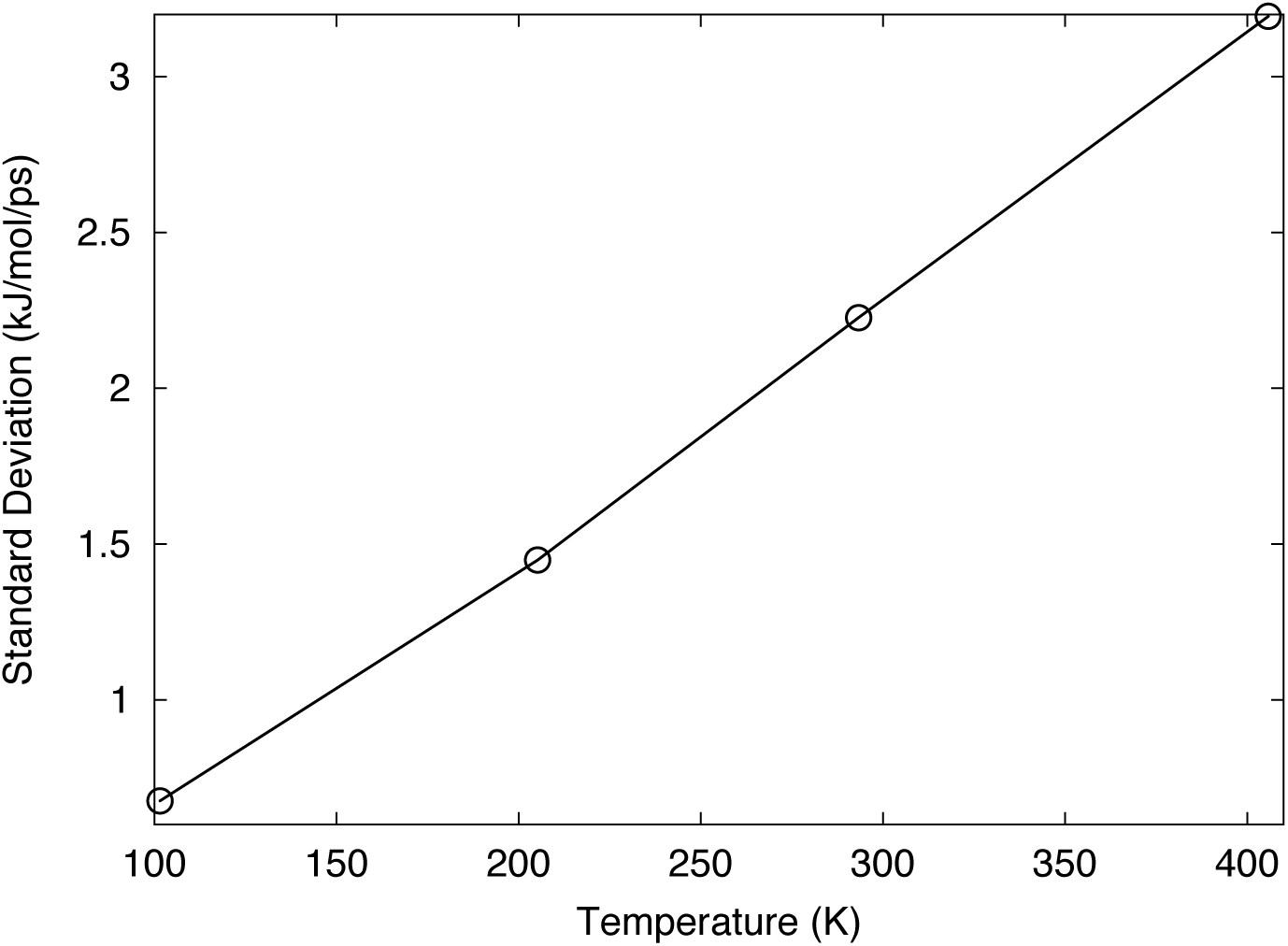
Standard deviation of the 1 ps energy change distribution versus temperature.

The above conclusions only apply to integration error. As we saw in section 4, numerical error is negligible compared to integration error at this step size. There can be many sources of numerical error, and different ones may scale differently with temperature. It is impossible to make any universal statements about how the energy drift will scale in situations where numerical error is significant.

## 6. Scaling With System Size

Next we consider how energy drift scales with the number of degrees of freedom in the system. To test this, we simulated two additional proteins: the villin headpiece[15] (582 atoms) and α-spectrin[16] (5078 atoms). All simulations used mixed precision and a 1 fs time step. No constraints were used, so the number of degrees of freedom is simply three times the number of atoms. The results are shown in Table 4.

**Table 4.** Statistics for simulations of proteins of varying sizes. The mean and standard deviation describe the distribution of energy changes over 1 ps intervals. The slope is from a linear regression to the curve of energy versus time for the whole simulation. All simulations used mixed precision, a 1 fs time step, and 5 ns simulation length.

| Protein | Atoms | 1 ps Energy Change Distribution |  | Slope<br>(kJ/(mol·ps)) |
| --- | --- | --- | --- | --- |
|  |  | Mean<br>(kJ/(mol·ps)) | Standard Deviation<br>(kJ/(mol·ps)) |  |
| villin | 582 | -0.000828 | 1.50 | 0.000116 |
| ubiquitin | 1231 | 0.00144 | 2.23 | 0.00119 |
| α-spectrin | 5078 | 0.0154 | 4.67 | 0.0179 |

The standard deviation shows a very clear O(√*n*) dependence on the system size: dividing it by the square root of the number of atoms gives 0.062, 0.064, and 0.066 for villin, ubiquitin, and ?-spectrin respectively. Unsurprisingly, values that are uncorrelated between degrees of freedom also are uncorrelated between time steps. At least on short time scales, the assumption that energy drift is proportional to system size clearly is incorrect.

The systematic drift is harder to evaluate, given the larger uncertainties in it, but it clearly grows faster than the random drift. If anything, it appears to be growing quadratically in the system size: dividing the slope by the square of the number of atoms yields 3.4·10^−10^, 7.8·10^−10^, and 6.9·10^−10^ for villin, ubiquitin, and ?-spectrin respectively. Possibly this reflects the fact that no cutoffs were used in these simulations, so the total number of interactions grows as the square of the number of atoms. If so, the scaling might be different when simulating a periodic system with Particle Mesh Ewald, but it would still presumably be at least linear.

A consequence is that the relative magnitudes of random and systematic drift depend on system size, and for a sufficiently large system the energy drift will be primarily linear even on short time scales. We emphasize again that this does not make systematic error “more important” than random error. Both types of error indicate that the equations of motion are not being integrated accurately. Just because errors produce energy changes that are uncorrelated between time steps or degrees of freedom, that does not mean they cannot still distort one’s results.

## 7. Accuracy and Energy Drift

We now turn to a fundamental question: is energy conservation actually a good way to assess the accuracy of a simulation? The total energy is merely one degree of freedom out of thousands in the system. It is, to be sure, a very important degree of freedom, but so are many others. If a simulation produces little error in the total energy, can we safely assume that it also produces little error in all the other degrees of freedom? Conversely, if a simulation is accurate (for some reasonable definition of the word), to what extent does that guarantee precise energy conservation over long time periods?

The Verlet algorithm is *symplectic*, which means that it very precisely conserves a modified or “shadow” Hamiltonian.[17] The shadow Hamiltonian is different from the true Hamiltonian of the system, but the two are closely related, so precise conservation of one generally implies reasonable conservation of the other as well. The upshot is that symplectic integrators have much better long term energy conservation than would be expected based on the integration error.

This provides an answer to our first question: long term energy conservation does *not* necessarily imply accuracy. On the contrary, it is virtually guaranteed that the integration error in most degrees of freedom is larger (possibly much larger) than that in the total energy. This makes energy conservation a particularly bad way of estimating accuracy. This applies only to integration error. As we saw earlier, a symplectic integrator does not reduce the energy drift from numerical error. Therefore, if a simulation has little long term energy drift, it is plausible to conclude that there is minimal numerical error in other degrees of freedom. Even this conclusion needs to be qualified, however. Long term energy drift is determined primarily by systematic error, not random error. Therefore the only really reliable conclusion one can draw is that, if there is significant numerical error in the simulation, it is of a type that only produces random drift in the energy, not systematic drift.

Furthermore, there can be many different sources of error in a simulation, each of which contributes to both the random and systematic energy drifts. There is no reason to expect their contributions to random drift to be correlated, but by definition, all sources of systematic drift are correlated. If some sources produce a positive systematic drift and others produce a negative systematic drift, they will tend to cancel out, leading to an artificially low drift on long time scales. Increasing the error can actually decrease the energy drift and make the simulation appear more accurate than it really is.

To illustrate this effect, we performed a series of 5 ns simulations of ubiquitin in which the SHAKE algorithm[18] was used to constrain the lengths of bonds involving hydrogen to their equilibrium values. This is a common technique to eliminate fast motions and allow a larger time step to be used. It introduces a new source of error, however: constraints are implemented with an iterative algorithm, so the constrained distances only remain constant to within a user specified tolerance. Table 5 shows the results of three simulations with different combinations of step size and constraint error tolerance.

**Table 5.** The cancelling effects of step size and constraint error tolerance on energy drift. The drift values are calculated from a linear regression to the curve of energy versus time for the whole simulation. All simulations were 5 ns long and used mixed precision.

| Step Size (fs) | Constraint Tolerance | Energy Drift<br>(kJ/(mol·ps)) |
| --- | --- | --- |
| 2.0 | $10^{-6}$ | 0.00424 |
| 2.0 | $10^{-4}$ | -0.950 |
| 2.6 | $10^{-4}$ | -0.0201 |

The first simulation uses a 2 fs step size and constraint error tolerance of 10^−6^. These are commonly used values, and produce a reasonably small energy drift. In the second simulation, we increase the constraint error tolerance to 10-4. This introduces large errors into the simulation, and causes the energy to decrease rapidly. In the final simulation, we increase the step size to 2.6 fs. This introduces additional large errors into the simulation and, on its own, would cause a rapid increase in energy. But because the two sources of error have opposite effects on systematic energy drift, they mostly cancel out and the actual energy drift is much smaller than in the previous case. That does not mean they have no effect, of course. This is a very inaccurate simulation, and no results calculated from it should be trusted. But it is impossible to discover that inaccuracy just by looking at long term energy drift.

What about the second question: to what extent does accurate integration guarantee long term energy conservation? The true equations of motion conserve energy exactly, so a perfect integrator would of course do so as well. In practice, no integrator is perfect so we should expect errors in all degrees of freedom, including the total energy. A symplectic integrator makes much smaller errors in the energy than in other degrees of freedom. This is usually presented as an advantage: given the same level of error, a symplectic integrator will conserve energy better than a non-symplectic one. This comparison can easily be reversed, however: given the same level of energy conservation, a non-symplectic integrator will be more accurate than the symplectic one. It is thus entirely possible for one integrator to be more accurate, but a different one to conserve energy better.

There has been much research into integration methods over the years, and many of the most widely used ones are non-symplectic.[19] This includes, for example, the very popular Runge-Kutta and Adams-Bashforth families of integrators. Much of the research in recent decades has involved error-controlled, variable time step integrators that continuously adjust the step size (and sometimes integration order) to keep the total error below a user-specified limit.[20] This allows them to be significantly more efficient than fixed step size integrators. It also makes them robust against rare events when, for example, two atoms come unusually close to each other. Events of this sort have been shown to be a source of error in molecular simulations[17]; variable time step integrators eliminate these problems by simply detecting the close collision and reducing the step size for a few time steps. For these reasons, they are now widely used in such fields as mechanical engineering, aerospace, and fluid dynamics.

The value of error controlled integrators can be illustrated with a simple example suggested by So? derlind.[20] We simulate a single charged particle moving in an elliptical orbit around a fixed charge. The trajectory generated with a fixed step size Verlet integrator is shown in the top half of Figure 6. Accumulated error causes the orbit to precess with time. This has no effect on the energy of the system, which is well conserved over the simulation, yet the particle still clearly ends up in the wrong location.

**Figure 6.**
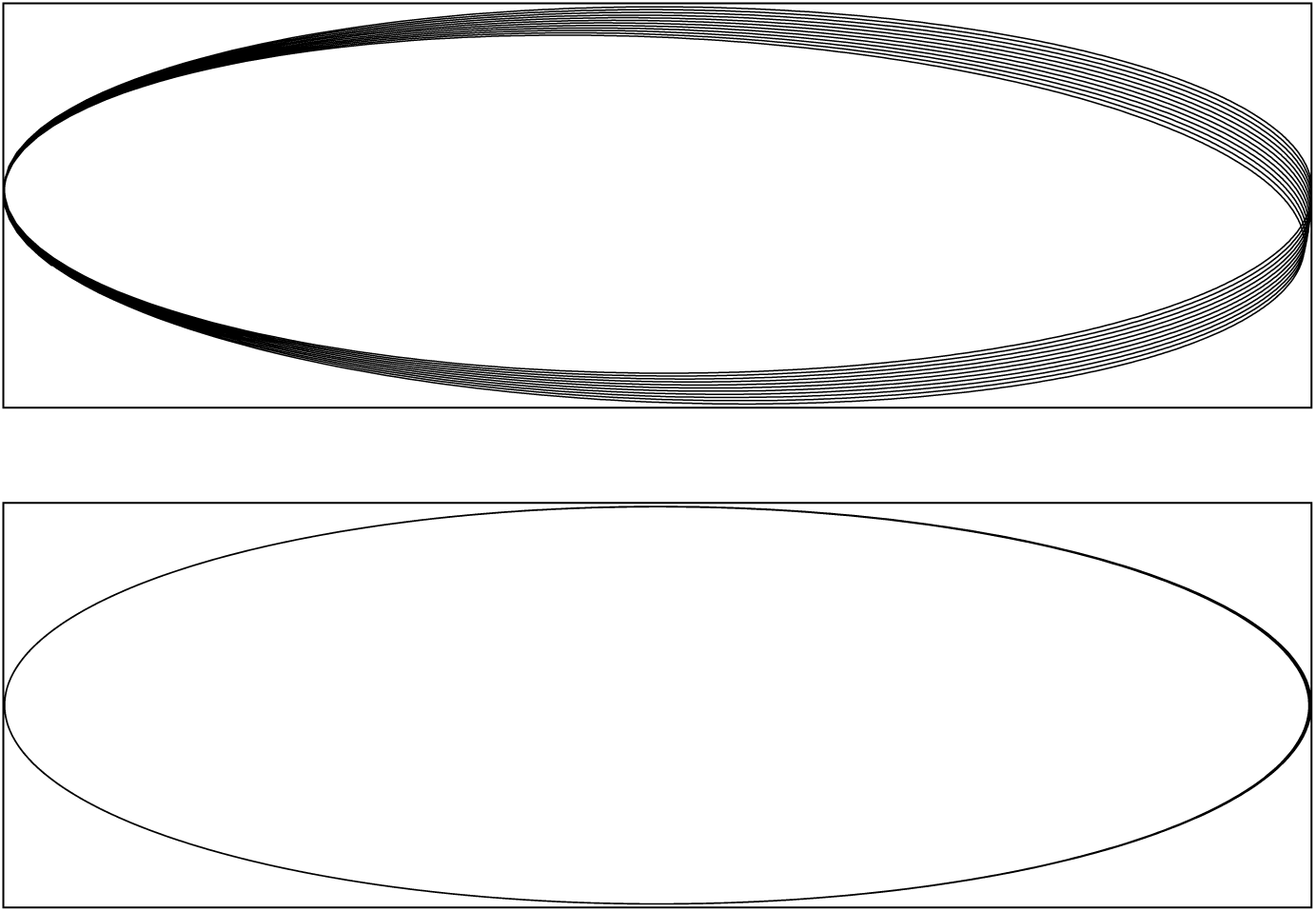
The trajectory of a particle in an elliptical orbit as integrated with a fixed time step (top) or variable time step (bottom) integrator. The average step size was the same in both cases.

OpenMM also provides an error controlled variable time step Verlet integrator. We simulated the same system, adjusting the error tolerance so the average step size over the simulation was identical to the previous case. The trajectory is shown in the bottom half of Figure 6. This time, there is negligible precession. The total number of time steps is identical, but the integrator is able to concentrate them in the parts of the trajectory where they are most needed, leading to a substantially more accurate result at similar cost. This is true despite the fact that the variable time step integrator is not symplectic.

This example is much simpler than most molecular simulations, and it may not be representative of them. Further research is clearly needed to understand how well the variable step size integrator reproduces statistical properties. Still, this example is valuable for providing a clear graphical illustration of how energy conservation does not imply accuracy. The fixed step size integrator produces an objectively less accurate result than the variable step size one, despite the fact that it is symplectic and has less energy drift over long time scales. (The energy drift over this short time interval was negligible for both simulations.)

It may be time to reconsider the molecular simulation community’s reliance on fixed step size Verlet integrators. More modern algorithms have many advantages in terms of both robustness and efficiency. Many of them are not symplectic, which means they will appear inferior as long as one only looks at the long term drift in the total energy, but this may be nothing more than an artifact of ignoring most types of error in the simulation.

## 8. Conclusions

We have performed simulations to test three commonly made assumptions about energy drift in molecular simulations: that it is linearly proportional to simulation length, linearly proportional to temperature, and linearly proportional to the number of degrees of freedom. We find that all three of these assumptions are wrong. This means that most published numbers for energy drift must be treated with extreme caution. Values computed for different systems or with different simulation parameters are not directly comparable to each other.

We also find that energy drift is too complicated to be accurately described with just a single number. A more thorough analysis is required to distinguish between random drift versus systematic drift, and between integration error versus numerical error. For example, by building a histogram of energy changes over short time intervals, one can separate systematic drift from random drift and measure each one independently. By measuring how they change as the step size is varied, one can then determine whether they are primarily caused by integration error or numerical error. These different factors may scale in very different ways with step size, simulation length, system size, and other simulation parameters. All of them are important and they need to be measured independently.

We further argue that energy conservation is not a reliable way to assess the accuracy of a simulation. For example, multiple sources of drift may cancel to produce good energy conservation even for an inaccurate simulation, as shown in Table 5. Conversely, a symplectic integrator can be objectively less accurate than a non-symplectic one, as shown in Figure 6, even though it does a better job of conserving energy over long time scales. In fact, when using a symplectic integrator such as fixed time step Verlet, it is virtually guaranteed that the integration errors in most degrees of freedom are larger than those in the total energy. This makes energy conservation a uniquely bad proxy for accuracy. At best, it is a necessary but not sufficient condition to conclude that one’s results are accurate.

## 9. Acknowledgments

The authors thank John Chodera and Michael Sherman for useful discussions and comments during the development of this manuscript. This work was supported by Simbios via the NIH Roadmap for Medical Research Grant U54 GM072970.

